# Skip segment Hirschsprung disease: modelling the trans-mesenteric origin of the enteric nervous system in the human colon

**DOI:** 10.1101/2019.12.22.886606

**Authors:** Donald F Newgreen, James M Osborne, Dongcheng Zhang

**Author notes:** Correspondence to: Donald F Newgreen, Murdoch Children’s Research Institute, Royal Children’s Hospital, Parkville 3052, Victoria, Australia. Contributions: Concept DFN; acquisition and analysis of data: DFN, DCZ; mathematical modelling JMO; drafting of the manuscript DFN; editing/revision of the manuscript DFN, DCZ, JMO.

## Abstract

Skip segment Hirschsprung disease is a difficult to explain human enteric neuropathy where a ganglionated region lies within a region of total colonic aganglionosis.. Recently, trans-mesenteric migration was described in the mouse intestine whereby neural crest cells migrate via the mesentery across a U-shape gut loop from the midgut to the hindgut: this could explain skip segment Hirschsprung disease. To investigate this, human intestinal growth parameters were derived from published sources and correlated with enteric neural crest cell migration. These processes were then simulated using agent based mathematical models scaled to human intestinal growth. A Hirschsprung-associated slowing of migration was imposed and trans-mesenteric migration was allowed. From the developmental anatomy we conclude that trans-mesenteric migration is unlikely in normal human embryogenesis, but with a Hirschsprung-associated slowing of enteric neural crest cell migration it could occur at Carnegie stages 17 and 18. By varying the division rate of enteric neural crest agents we could reproduce full colonisation, short segment, long segment and skip segment Hirschsprung and hypoganglionic segments.

**Summary Statement:** Skip segment Hirschsprung disease in humans challenges current explanations. Mathematical modelling shows how this birth defect could develop.

## INTRODUCTION

Hirschsprung Disease (HSCR) is a long-known (Sergi, 2015) and relatively common birth defect (East Asian: 1/3200, Caucasian: 1/5000 live births) in which the distal-most intestine lacks neurons of the enteric nervous system (ENS)(Heuckeroth, 2018; Tilghman et al., 2019). ENS neurons aggregate into numerous ganglia, hence HSCR is also known as aganglionosis (McKeown et al., 2013). The aganglionosis in about 80% of cases is confined to the rectosigmoid colon and is termed ‘short segment’, but ‘long segment’ variants occur, including total colonic agagnglionosis (TCA). The result of this is an inability to pass intestinal contents through the aganglionic segment; this produces a distension proximal to the affected region graphically termed megacolon. Current treatment is identification of the aganglionic region by biopsy (Kapur and Kennedy, 2013), surgical resection of this segment plus a safety margin into the more rostral ganglionated zone, followed by colostomy or, if possible, anastomosis of the proximal gut segment to the rectum. Without this treatment the patients suffer bouts of potentially fatal enterocolitis (Pontarelli et al., 2013), but morbidity is a problem even with surgery (Dickie et al., 2014; Rintala and Pakarinen, 2012).

The use of avian (Goldstein and Nagy, 2008) and rodent (Zimmer and Puri, 2015) models has led to an understanding of the embryological origin of HSCR. Most ENS cells are derived from vagal-level (ie. hindbrain) enteric neural crest (ENC) cells (Hutchins et al., 2018; Simkin et al., 2019). These cells migrate into the oral end of the embryonic gastrointestinal tract and colonize the rest of the gut to the rectum in an oral-to-anal wave of cell migration in the gut mesoderm (Young et al., 1998). A slowing or premature stoppage of this migration wave, amplified by the simultaneous growth of the gut domain within which ENC cell colonisation is occurring, explains why the segment of aganglionosis in HSCR is the distal-most segment (Landman et al., 2007). The identification of additional independent sources of ENS cells from the sacral NC (Burns and Le Douarin, 1998) and mid-trunk NC (via Schwann cell precursors) (Uesaka et al., 2015) has not led to fundamental changes to this sequence because these sources provide fewer ENS cells compared to the vagal ENC-derived cells, and cannot compensate for lack of the vagal ENC cells (Burns et al., 2000; Hearn and Newgreen, 2000). Observations of ENS formation in human embryos (Attie-Bitach et al., 1998; Fu et al., 2003; Fu et al., 2004; Wallace and Burns, 2005) are consistent with the view that the human ENS develops similarly to that in the animal models.

Despite this emphasis on the distal location of aganglionosis, a rare variant termed skip segment HSCR also occurs (Martin et al., 1979). In this form, an ‘island’ of ganglionated bowel of variable size and variable ganglion density occurs with aganglionic segments both oral and anal to it (**Fig. 1**). Combining the literature reviewed by O’ Donnell and Puri (O’Donnell and Puri, 2010), Ruiz et al. (Ruiz et al., 2016) and Coe et al. (Coe et al., 2016), there have been over 30 patients with skip-like ENS defects described since 1954. Most (>90%) skip segments have been detected in a TCA region (O’Donnell and Puri, 2010). Skip segments are typically in the caecum, the ascending colon or the transverse colon (Fig. 1). Skip segments in the more distal colon are rare (Alfawaz et al., 2017).

**Fig. 1.**
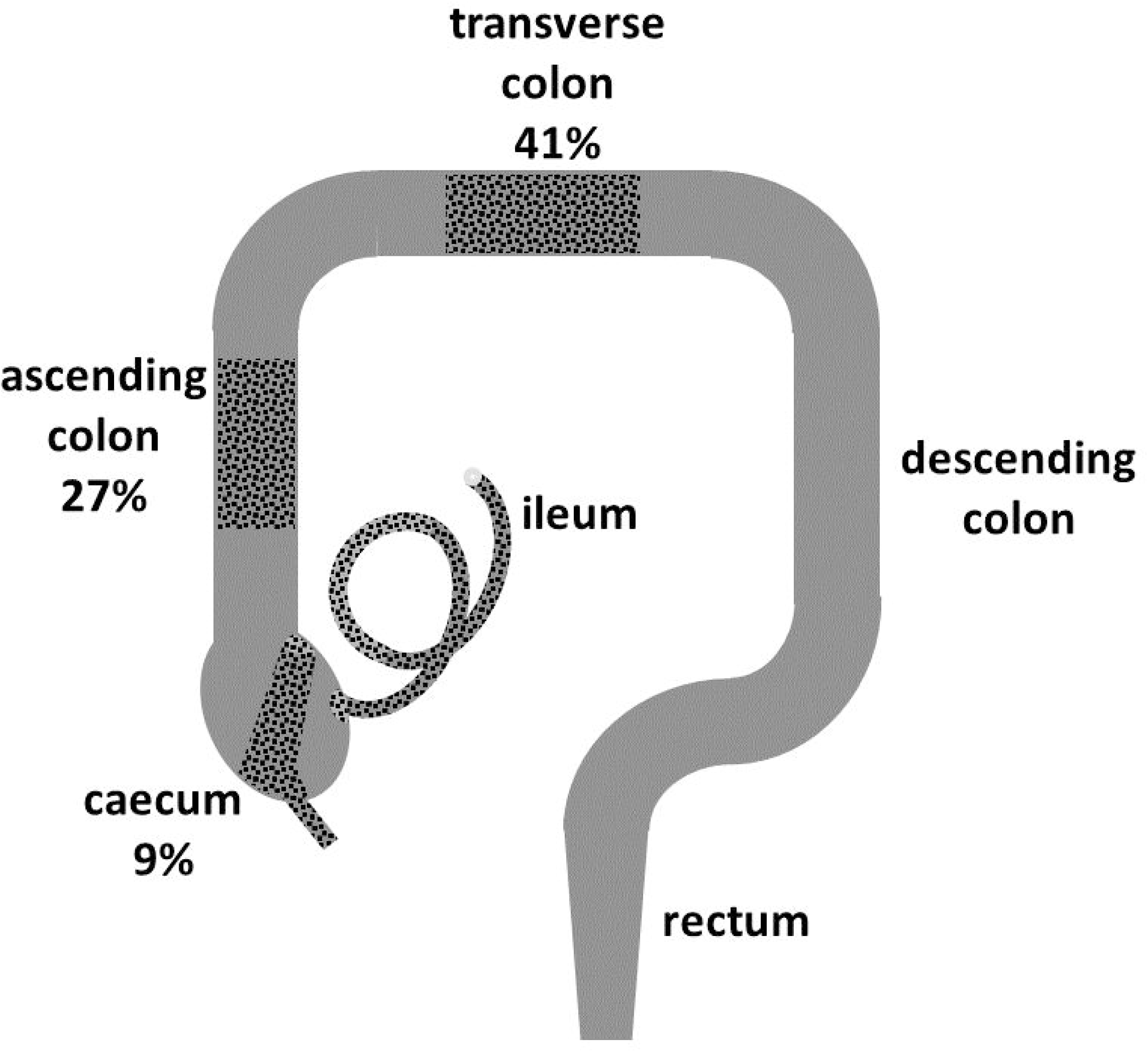
Locations and frequencies of skip segments in human large bowel. Ganglionated regions are indicated by black dots. The diagram is not indicative of the area of the ganglionated regions nor of ganglion cell density. From O’Donnell and Puri, 2010.

Conventional HSCR and skip segment HSCR are related genetically. The gene *RET* is the most commonly mutated gene in HSCR (Emison et al., 2010), and examination of *Ret51(C618F)* knock-in mice has revealed a post-caecal skip segment although most of these mice showed a simple TCA (Nishiyama et al., 2012). In addition, in humans, skip segments in a TCA context have been found in Waardenburg-Shah syndrome patients with homozygous mutation of *EDNRB* (Gross et al., 2015) and with hemizygous mutation of *SOX10* (Ruiz et al., 2016), both known HSCR genes. Nevertheless, skip segment HSCR obviously poses a serious problem for a developmental explanation because this lesion is not accommodated by a simple curtailment of the oral-to-anal wave of ENC cell migration.

In a pioneering discussion combining evidence from wild-type and *ls/ls* HSCR model mice and human clinical reports, Kapur and co-workers (Kapur et al., 1995) pointed out that ENC colonisation in the gut wall was delayed in the caecum of the mouse, and a stream of ENC-derived cells extended further distally extramurally at the border of the caecum and the mesentery. After by-passing the caecum, some of these cells appeared to re-enter the colon wall distal to the caecum. This gave a transient uncolonised gap proximally including the caecal region, as well as the distal uncolonised zone. Kapur et al suggested that normally the ENC cells ‘back-fill’ this gap, but failure to accomplish this could explain caecal and juxta-caecal skip segment HSCR in humans. However this explanation is not consistent with the 40% of skip segments located in the transverse colon.

More recently another explanation has arisen (Nishiyama et al., 2012) from time-lapse observations using advanced cell imaging of ENC cell movement in mouse embryo gut in organ culture, and in static observations of mouse and human gut. At a crucial stage of intestinal colonisation, the intestine forms a U-shaped bend with the proximal midgut as one arm, joined by a short span of mesentery to the arm consisting of distal midgut (including caecum) and hindgut. As intramural ENC cells colonise the proximal arm, individual ENC cells can use the mesentery as a bridge to wander across and enter the nearby distal arm. This short-cut places these ‘trans-mesenteric’ ENC cells well ahead of the intramurally colonising cells (termed ‘circumflex’ ENC cells (Nishiyama et al., 2012)). Due to the ‘frontal expansion’ mode of colonisation (Simpson et al., 2007), these trans-mesenteric ENC cells proliferate, taking over the colonisation of more distal gut. Assuming this trans-mesenteric event occurs *in vivo* as it does in organ cultures, then the stage at which this can occur is probably brief *in vivo* since further growth and morphogenesis increases the separation of the proximal and distal intestinal arms. This and other models have been incorporated in recent reviews (Coe et al., 2016; Sehgal, 2018), and this notion of Nishiyama and colleagues (Nishiyama et al., 2012) explains the occurrence and variable position of all skip segments better than the caecal by-pass model of Kapur and colleagues (Kapur et al., 1995).

Here we ask whether this concept of trans-mesenteric origin of ENS can be applied to both normal colonic ENS formation and skip segment HSCR in humans. Direct observation and experimentation are not possible in human embryos. Therefore from on-line and published sources we derive anatomical parameters of the forming intestine and ENS in human embryos in terms of rate of development, scale and morphology. We then use computer models of ENC cell colonisation dynamics that were previously successful in animal applications (Landman et al., 2014; Landman et al., 2007; Newgreen et al., 2013) but with human temporal and anatomical parameters. From these we attempt to define the likely effect of altering the timing, location, division rate and number of the trans-mesenteric ENC cells on the ENS phenotype and the development of skip segment HSCR. Our conclusions are that trans-mesenteric migration is unlikely in normal colonisation of the human distal intestine by ENC cells, but that with a HSCR-related slowing of migration, this mechanism explains the existence and most frequent sites of skip segment disease.

## RESULTS AND DISCUSSION

### Likelihood and location of trans-mesenteric ENC cell migration in human embryos

#### Gut morphogenesis and ENC cell colonisation in human embryos: lessons from the mouse and chick

The human embryonic mid- and hindgut undergoes extensive growth and looping (Fig. 2), as does the gut of the mouse and chick embryo. In the normal mouse, ENC cells migrate along the descending (proximal) intestinal loop at a stage when this is separated from the ENC-free ascending (distal) loop by a narrow mesentery, allowing trans-mesenteric colonisation (Nishiyama et al., 2012). By contrast, in the avian embryo, ENC cell advance occurs relatively early compared to intestinal looping (Allan and Newgreen, 1980; Southwell, 2006; Zhang et al., 2019). Uncolonised intestine is crucial for a new site of entry to become established because experiments in animal models and in mathematical models indicate that a resident ENS population suppresses the colonising ability of ENC cells arriving later (Meijers et al., 1992; Simpson et al., 2007). Thus trans-mesenteric colonisation by ENC cells does not occur in the looped chick embryo intestine because the distal intestine is already colonised by the oral-to-anal ENC cell wave.

**Fig. 2.**
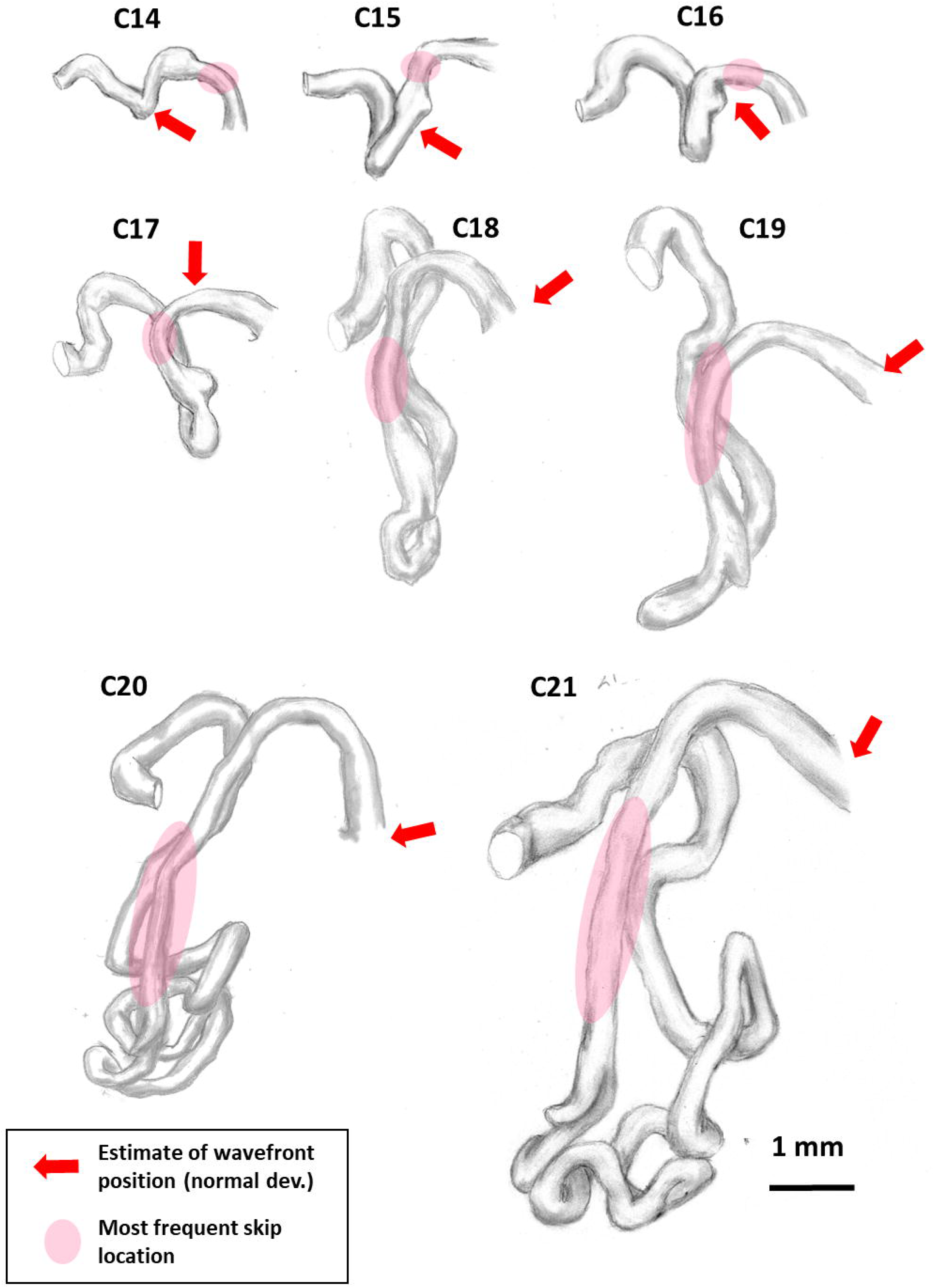
Sketches of intestine morphogenesis of the human embryo. These drawings are based on sections (VHE), reconstructions (3DHE) and wholemount images (THEF). The red arrow indicates the front of the ENC cell colonisation wave in normal development, estimated from Attie-Bitach *et al.* 1998; Fu et al., 2003, 2004; Wallace and Burns 2005; Nishiyama et al., 2012. The pink overlay area represents the trans-mesenteric insertion site in the post-caecal colon.

In the developing human, like the chick, the progress of the ENC cell wavefront (see Suppl. Fig. 8 of Nishiyama et al., 2012) has advanced along the intestine before the formation of the intestinal U-shaped loop that will bring the proximal midgut arm close to the caecal and post-caecal regions of the distal intestine (Fig. 2). Thus when the separation distance makes ENC cell migration across the mesentery feasible at the sites of skip segments occur (Fig. 1), the trans-mesenteric cells would likely encounter intestine populated with ENS cells that have migrated oro-anally along the intestine. We therefore conclude that in ENS morphogenesis in the normal human embryo, trans-mesenteric occupation of the more distal colon is unlikely. This does not mean that human ENC cells are intrinsically incapable of this movement, rather in normal development they lack the temporal opportunity.

#### ENC cell colonisation in human HSCR embryos compared to rodent models

In rat and mouse embryos with a distal HSCR-like aganglionosis, the progress of the ENC migratory wave is slowed proximally even in the region that is eventually colonised (Newgreen and Hartley, 1995; Shin et al., 1999). Similar generalised slowing is likely in affected human embryos, and in this case a latent capacity of ENC cells to migrate trans-mesenterically may be unmasked.

#### Spatio-temporal window for trans-mesenteric migration in human embryos

The developmental anatomy of the human gut at the most common sites of skip segments offers clues as to when and where trans-mesenteric migration might occur. Note that we assume the intestinal morphology in human HSCR embryos is normal and only the progress of the ENC cell migratory wave is affected.

In the development of the mouse intestine, the caecum is relatively close to the umbilical tip of the intestinal loop whereas in the human embryonic intestine at C14-16 the caecum is positioned relatively dorsal in the ascending limb of the intestinal loop (Fig. 2). This means that in human embryos, trans-mesenteric crossing by ENC cells before C16 would, at the region of least separation, lodge ENC cells oral to the caecum. Since skip segments do not occur here (see Fig. 1)(O’Donnell and Puri, 2010), trans-mesenteric crossing resulting in a skip would be unlikely to occur before C16. On the other hand, due to growth of the intestine at and after C19 (Fig. 2, 3) the separation increases and the mesentery becomes more tortuously folded which must hinder trans-mesenteric crossing. At C17 and C18 the more proximal descending intestine is adjacent to the proximal part of the post-caecal gut (Fig. 2). Consistent with this, most skip segments have been reported in this region rather than in the more distal post-caecal intestine (Fig. 1). At this embryonic stage the distance of closest approach measured directly across the mesentery is shortest (range 300-400 μm).

**Fig. 3.**
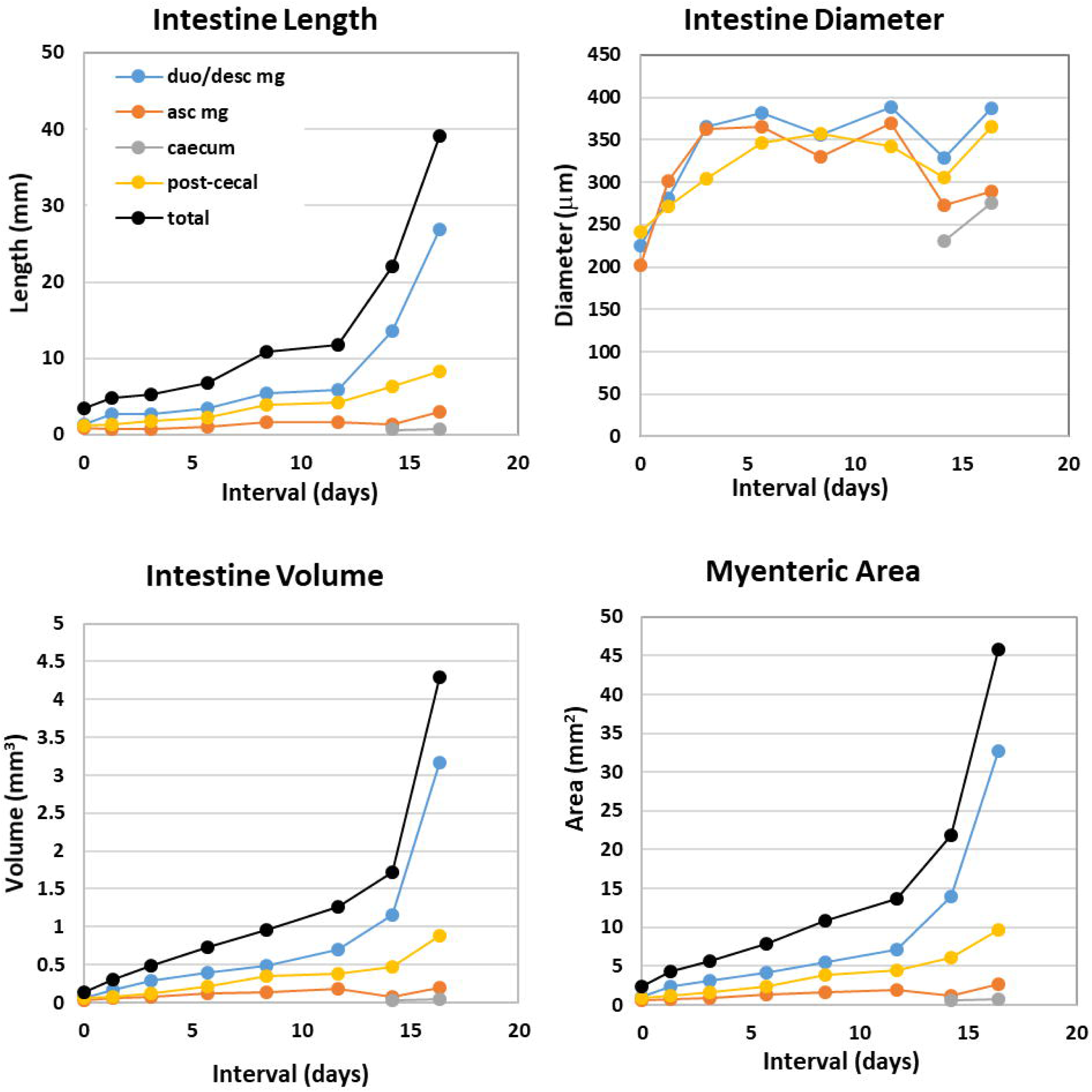
Human embryo intestinal growth parameters from C14 to C21. C14 (about 34 days post-fertilization) is set at 0 days and C21 (about 50 days post-fertilization) is set at about 16 days. Key: Gut from bile duct and midgut to tip of loop= duo/desc mg. Midgut from tip of loop to mid-caecum= asc mg. Distal gut from caecum to cloaca=post-caecal. Tubular caecum (after C20)=caecum.

We therefore conclude that, with a slowed ENC colonisation wave, the most likely time for trans-mesenteric crossing would be stages C17/C18. The length of closest approach of the proximal and distal limbs of the intestinal loop, and therefore most likely position of crossing, extends for about 1 mm along the caecal and post-caecal intestine at C17/C18. The occurrence of a temporal and spatial window for trans-mesenteric crossing has already been described in the mouse intestine (Nishiyama et al., 2012).

### CA models of ENC cell colonisation reproduce conventional and skip segment HSCR

#### Intestinal growth measurements in human embryos

The increase with developmental time in the length, diameter and volume of various segments of the human embryonic intestine (assuming that the gut is approximately circular in local cross-section at each point) from C14 to C21 (age post-fertilization about 34 to 50 days) is given in Fig. 3. The time intervals between stages are set with stage C14 as starting time (nominal t=0 day). The area of the potential myenteric plexus is also given because this is the layer occupied by ENC cells migrating and proliferating within the gut mesoderm. This is the correlate of the 2D surface approximated in grid-format by the CA model described previously. Noteworthy is the rapid growth of the proximal (descending) midgut compared to more distal intestine, which is also observed in other species (Newgreen et al., 1996). These data are used to scale the CA model of ENC colonisation.

#### Skip segment HSCR can emerge in CA models of ENC cell colonisation

ENC cell number and division indirectly drives net directional motility (Barlow et al., 2008; Simpson et al., 2006; Simpson et al., 2007). HSCR mutations (Amiel et al., 2008) in ENC cells reduce their division and migration (Gianino et al., 2003; Hearn et al., 1998; Nagy and Goldstein, 2006; Wu et al., 1999; Young et al., 2001). This leads to failure of colonisation distally (ie. HSCR) if migration fails to exceed intestinal myenteric layer expansion (Newgreen et al., 1996). Previous CA models have explored a range of rates of division of ENC agents, with the focus being to produce ENC agent colonisation that ‘just failed’ (ie. short segment HSCR) or ‘just succeeded’ (Binder et al., 2015). However skip lesions occur most frequently in a TCA context (Coe et al., 2016; O’Donnell and Puri, 2010), which is a major failure of colonisation. This suggests a greater reduction in ENC division rate is at play.

For the CA model we derived exponential growth curves to represent the rate of human intestinal expansion (Fig. 4A). If trans-mesenteric crossing was disallowed, then HSCR of variable length could evolve depending on the values of agent movement P_M_ and agent division P_D_ (Suppl. Fig. S1). Then to model the ENC cell migration process (Fig. 5A, B), we imposed (Fig. 5D) a transfer of ENC agents from the descending midgut to the post-caecal gut in the spatial context of the C17/C18 stage (choice of this stage is discussed above). This was achieved by allowing cells within a specified spatial and temporal window (red in Fig. 5D) to move to sites in a region of post-caecal intestine. Once transferred to the distal arm, the agents proliferated and moved with the same parameters as agents in the originating proximal population. Note that ENS-free space in the new site is available in all directions and the CA modelled ENC agent population responds by expanding in all directions. This is likely to occur biologically because ENC cell migration directed by the availability of ENS-free space has been demonstrated experimentally in the mouse and avian intestine (Anderson et al., 2007; Burns et al., 2002; Nishiyama et al., 2012; Simpson et al., 2007). Under these conditions a skip segment may be generated and may persist (Fig. 4B).

**Fig. 4.**
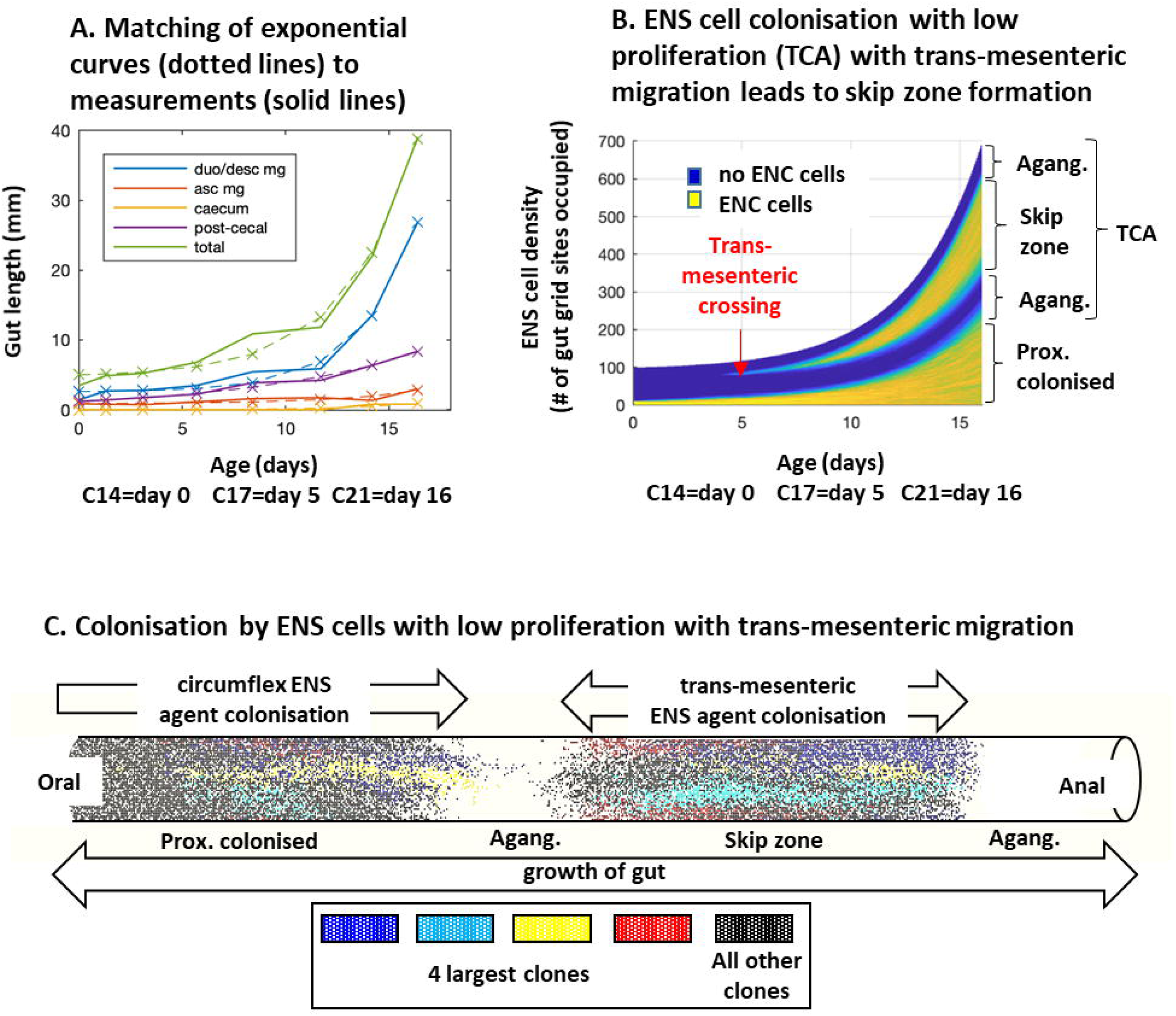
Modelling of skip segment formation. **A**. The growth parameters of the intestine can be approximated accurately by exponential curves. **B.** Single simulation of the time course of colonisation of the growing intestine with low rate of division P_D_ to cause TCA with superimposed trans-mesenteric crossing. Here the ENC agent P_D_ and the rate of intestinal expansion are balanced such that the lengths of the aganglionic zones flanking the skip segment do not change. Developmental time=x-axis, gut length=y-axis. **C.** Plotting of ENC agent clones indicates dominance of a few clones (see Cheeseman et al., 2014) in both the proximal colonised segment and in the skip segment, but the relative contribution of clones is not identical in the two segments.

**Fig. 5.**
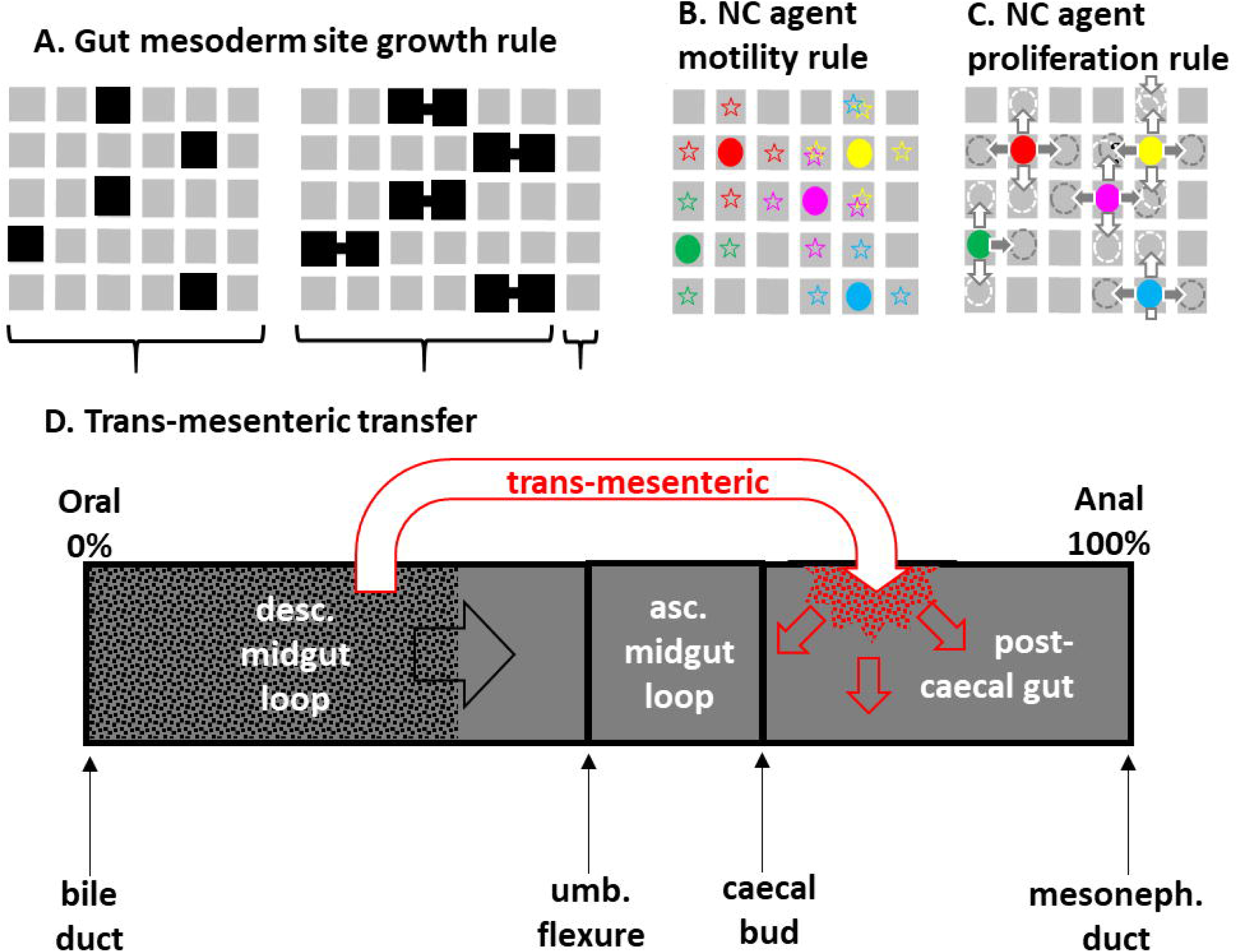
Scheme for modelling circumflex and trans-mesenteric ENC agent colonisation. **A.** The gut mesoderm layer (grey, gridded) increases in length at each timestep by choosing at random one grid site (black) in each row to duplicate, intercalating the new site into the row. **B.** At probability dtP_M_ each ENC agent (coloured circles) may move to any one of four adjacent sites (stars) chosen at random. Double occupancy is prevented by the second occupant’s move being aborted. **C.** At probability dtP_D_ each ENC agent (coloured circles) may divide, the two daughter agents (dotted circles) being placed randomly (arrows) north/south or east/west of the original position. Double occupancy is prevented by the second occupant’s division being aborted. **D.** Concept of CA model of severe HSCR (TCA) with trans-mesenteric crossing. Gut myenteric layer is shown in dark grey. This gut layer expands in length according to Fig. 5A. Circumflex ENS agents (black dots) colonize oro-anally in this gut layer (black arrow). Conditions of ENC agent division and migration versus gut expansion are used such that these ENC agents do not colonise past the caecum, equivalent to TCA. Some ENC agents are translocated (in red) from descending midgut to distal to caecum, by including post caecal sites in the neighbours of the selected cells, to model the trans-mesenteric migration process. These agents then colonise the post-caecal gut in all directions (red arrows) with the same division and motility parameters as the proximal ENC agents.

Normal ENC agent division during colonisation evolves such that a relatively few clones at early stages contribute disproportionately to the final ENS population (Cheeseman et al., 2014). The same type of clonal inequality occurs in the skip region in the model, but the precise identities and proportions of the clonal cells in the skip segment may not be identical to that in the distal circumflex ENC population (Fig. 4C).

#### Varying the division of ENC agents in CA models can predict all forms of HSCR

Depending on the setting of the probability of ENC agent division *P*_D_, outcomes emerged in the models representing the normal fully colonised ENS, conventional HSCR (distal aganglionosis), skip segment HSCR and hypoganglionic skip segment (Suppl. Fig. S1-3).

When a skip emerged because, for example, the ENC agent division setting *P*_D_ was low to ensure a TCA context and because the distal part of the colon was not involved in the U-shaped intestinal loop, the skip segment was always in the proximal part of the post-caecal segment, in agreement with real site of skip segments (Fig. 1). When trans-mesenteric crossing was imposed with the ENC agent rate of division set relatively high, proximal and distal aganglionic zones were transient, giving a fully colonised ENS (Suppl. Fig. S2-3). By decreasing the ENC agent division rate *P*_D_, a conventional non-skip HSCR developed via a transient skip segment (Suppl. Fig. S2,3). This was due to the population of ENS agents derived from the trans-mesenteric agents having time to expand orally far enough to fuse with the anally directed intramural ENC agent wave. In this case the distal-most ENS was almost entirely composed of trans-mesenteric daughter agents as in the mouse experimental model (Nishiyama et al., 2012). However the agents failed to complete colonisation distally because, due to their low division setting, they were out-distanced by the expansion of the grid representing the growing distal gut mesoderm. Further reduction of *P*_D_ generated skip segment HSCR (Fig. 4B, Suppl. Fig. S2), and if reduced still further the skip zone became progressively hypoganglionic to a point resembling TCA.

The timing/positon of the trans-mesenteric crossing can play an important role in the outcome of the developing gut. If the skip happens early enough it can allow the a variety of phenotypes including skip segment HSCR. On the otherhand if the crossing occurs later then the trans-mesenteric cells fail and the skip segment phenotype is transient if it appears at all (Suppl. Fig. S3). More complete presentations of the variability of outcome in these cases as P_M_ and P_D_ are varied is given in Suppl. Figs. S1-3.

## Conclusions

How skip segment HSCR is generated has been a puzzle, and the mechanism of formation of skip segment HSCR presented here arose from recent studies of ENS formation in the normal mouse embryo, where migration of ENC cells across a mesenteric bridge from midgut to hindgut was detected (Nishiyama et al., 2012). However descriptions of the morphogenesis of the intestine and ENC colonisation presented here suggests slight but important differences in the relative timing of these two events between human and mouse embryos. Our interpretation is that, relative to intestinal looping, the progress of ENC colonisation in the human is advanced over that in the mouse. This in turn suggests that, unlike in the mouse intestine, the trans-mesenteric mode may be of little or no importance for the normal ENC-colonising wave in the human.

Despite this conclusion in the normal case, the observations presented here suggests that an HSCR-like slowing of ENC colonisation produces conditions in human embryonic intestine that would allow trans-mesenteric migration. This trans-mesenteric mechanism (Nishiyama et al., 2012) and the caecum bypass model (Kapur et al., 1995) reproduce caecal and juxta-caecal skip sites, and both models also account for the observed dearth of skip segments in the distal colon. However, this trans-mesenteric model also accounts for the most frequent sites of the skip in post-caecal gut, corresponding to the future ascending and transverse colon. In addition, the CA model shows how simply varying one numerical parameter, the probability of division of ENC cells, can simulate the full spectrum of HSCR defects such as short-segment, skip segment, and long segment such as TCA, and also patchy and hypoganglionic segments.

We conclude that this trans-mesenteric model offers a convincing explanation for the existence, location and diversity of skip segment HSCR. We propose that skip segments are more frequent in the TCA region than is currently recognised, and this could be revealed by more closely sampling along its length (Kapur, 2016) using neuronal markers rather than conventional histology, to identify short skips, easily missed hypoganglionosis and circumferential patchy ENS distributions (Takahashi et al., 2013).

## MATERIALS AND METHODS

### Human embryo intestinal growth parameters

Gut measurements were derived from images in online data bases: the Virtual Human Embryo (VHE) (http://www.ehd.org/virtual-human-embryo/) (Gasser et al., 2014), 3D Atlas of Human Embryology (3DHE) (https://www.3dembryoatlas.com/) (de Bakker et al., 2016), the Multidimensional Human Embryo (http://embryo.soad.umich.edu/) and The Collection of Immunolabeled Transparent Human Embryos and Fetuses (THEF) (https://transparent-human-embryo.com/) (Belle et al., 2017). These were supplemented with published histological and immunohistological sectional figures of ENS development (Attie-Bitach et al., 1998; Fu et al., 2003; Fu et al., 2004; Wallace and Burns, 2005) of human ENS development, reported with weekly increments. Fixed specimens were supplied by Dr T. Attie-Bitach (see ref. (Nishiyama et al., 2012)). Stated chronological ages of human embryos vary with the source; we follow the post-fertilizational ages used by O’Rahilly and Müller (O’Rahilly and Muller, 2010), focussing on Carnegie stages C14 (33-35 days) to C21 (49-52 days). A reasonable estimate of age in days at the stages considered can be made by adding 29 to the Greatest Length (GL) in mm.

Measurements of intestinal diameter and length were made using the VHE measurement tool and with Fiji (https://imagej.nih.gov/ij/). The diameter of the future myenteric plexus layer eventually occupied by migrating ENC cells was approximated to the gut diameter, since migrating ENC cells in the midgut and hindgut are positioned close to the outer border of the intestine. This enabled the circumference to be calculated, assuming circularity in cross-section. The length multiplied by the circumference equates to the potential myenteric area that ENC cells colonise.

We made corrections to certain measurements from VHE. For example, the GL of the whole C14 embryo (VHE #6502) from summing the sections (stated thickness 10 μm) was almost twice that of both the imaged whole specimen and the C14 average GL in O’Rahilly and Müller (2010). Since the latter dimensions are likely to be correct, the simplest explanation for this discrepancy is that the section thickness was 5 μm. In section images of the C15 embryo (VHE #721) point-to-point distances derived from the online measurement tool did not match the dimensions of the same points in the whole image of the same embryo. We assumed that the scale of the imaged whole embryo was correct as it matched published dimensions for C15 embryos (O’Rahilly and Muller, 2010). We therefore introduced for this specimen an *ad hoc* correction factor of x1.85 for measurements from sections using the VHE imaging tool.

### General properties of mathematical model of ENC cell colonisation

The mathematical models are based on those developed previously (Binder et al., 2012; Zhang et al., 2010; Zhang et al., 2018) involving cellular automata (CA), as reviewed by Newgreen and Landman and colleagues (Landman et al., 2014; Newgreen et al., 2013). Briefly, colonisation of the mid and hindgut by ENC cells – termed ENC agents in the model - are based on occupation of the myenteric area. This is simulated in a 2D periodic rectangular grid representing the myenteric layer in the gut mesoderm. This 2D format was chosen since normal colonisation occurs in a narrow layer in the mesoderm corresponding to the future myenteric plexus. This 2D grid grows in length (x-axis) by cycles of intercalary insertion of more grid spaces (Fig. 5A). Importantly, in this case the rate of growth is scaled exponentially to the growth parameters of the human intestine which were measured (see Fig. 4A). Here the domain length is given by the formula *L = a* + *b e*^*ct*^ where parameters were chosen to fit the data in Fig. 4A (*a=4.7390, b=0.2678, and c=0.2954*).

The ENC agents occupy intestinal grid spaces, being initially distributed at the oral end. As a convention, this is the left or west end. In each timestep (of duration *dt*) each of these ENC agents can move to an adjacent grid space at a probability *dtP*_M_ (Fig. 5B, note we use this rather than *P*_M_ so we can change the timestep without changing the properties of the two). Grid adjacency is defined as those which share an edge (Fig. 5B) or those connected by a trans-mesenteric bridge (Fig. 5D). Each ENC agent can divide to produce two agents with a probability *dtP*_D_, the daughters being placed in adjacent grid spaces (Fig. 5C). Both the direction of ENC agent movement to a new grid space and the placement of daughter agents is stochastic as this resembles normal colonisation viewed in time-lapse (Young et al., 2014). An exclusion rule is imposed so that attempts by an ENC agent to move into, or place a daughter into, an occupied grid space results in that action being aborted for that cycle. Each grid space is 250 μm wide and represents approximately 25 cell diameters.

A timestep of *dt=0.01* is used and ENC agent movement of 0.01 grid spaces per time unit (*dtP*_m_ =0.01), approximating to 2.5 ⍰m per hour representing slower movement than the 10 ⍰m per ¼ hour, based on observations from Young and colleagues (Young et al., 2004) and Druckenbrod and Epstein (Druckenbrod and Epstein, 2005, 2007) for cells in the mouse embryos gut in organ culture, and for cells crossing trans-mesenterically (Nishiyama et al., 2012). Agent division probability *dtP*_D_ is varied from 0.001 to 0.1 in different simulations with low probabilities reflecting low rate of division and hence a long segment HSCR condition. In the Supplementary figures we show the full range of possibilities by varying the movement probability, *dtP*_M,_ from 0.001 to 0.1.

The crucial point of difference between these simulations and previous models is that here some ENC agents are permitted to exit proximally and re-insert more distally ahead of the intramural wavefront. The exit and re-insertion sites are described by percentage of the gut domain length, as shown in Fig. 5D. This represents trans-mesenteric migration as described previously in the mouse (Nishiyama et al., 2012). The skip is implemented by including distal post post-caecal sites in the possible neighbour set for selected descending intestine sites. Specifically, here we consider the following cases: no skip (Suppl. Fig. S1); early skip (exiting from 20-25% and re-inserted into 70-75% of the gut; Suppl. Fig. S2); and late skip (from 40-45% and re-inserted into 90%-95% of the gut; Suppl. Fig. S3). This tran-mesenteric site is active between chosen times (here taken as 4 to 10 hours to approximately match C17 to C18).

## ACKNOWLEDGEMENTS

We would like to thank Dr Raj Kapur for helpful discussions, and similarly Profs. Heather Young and Hideki Enomoto. We would also like to thank the constructors of the on-line human databases mentioned in the text, for their enormous energy and tenacity in producing these invaluable resources.

